# Controlling the SARS-CoV-2 outbreak, insights from large scale whole genome sequences generated across the world

**DOI:** 10.1101/2020.04.28.066977

**Authors:** Jody Phelan, Wouter Deelder, Daniel Ward, Susana Campino, Martin L. Hibberd, Taane G Clark

## Abstract

**Background:** SARS-CoV-2 most likely evolved from a bat beta-coronavirus and started infecting humans in December 2019. Since then it has rapidly infected people around the world, with more than 4.5 million confirmed cases by the middle of May 2020. Early genome sequencing of the virus has enabled the development of molecular diagnostics and the commencement of therapy and vaccine development. The analysis of the early sequences showed relatively few evolutionary selection pressures. However, with the rapid worldwide expansion into diverse human populations, significant genetic variations are becoming increasingly likely. The current limitations on social movement between countries also offers the opportunity for these viral variants to become distinct strains with potential implications for diagnostics, therapies and vaccines.

**Methods:** We used the current sequencing archives (NCBI and GISAID) to investigate 15,487 whole genomes, looking for evidence of strain diversification and selective pressure.

**Results:** We used 6,294 SNPs to build a phylogenetic tree of SARS-CoV-2 diversity and noted strong evidence for the existence of two major clades and six sub-clades, unevenly distributed across the world. We also noted that convergent evolution has potentially occurred across several locations in the genome, showing selection pressures, including on the spike glycoprotein where we noted a potentially critical mutation that could affect its binding to the ACE2 receptor. We also report on mutations that could prevent current molecular diagnostics from detecting some of the sub-clades.

**Conclusion:** The worldwide whole genome sequencing effort is revealing the challenge of developing SARS-CoV-2 containment tools suitable for everyone and the need for data to be continually evaluated to ensure accuracy in outbreak estimations.

## Background

Within the last 20 years, three ‘spill-over events’ have occurred where a coronavirus has entered the human population from an animal host and cause significant morbidity and mortality. In the most recent event, the currently emerging SARS-CoV-2 virus - associated with disease COVID-19 - has spread rapidly worldwide to infect over 4.9 million individuals and has been associated with more than 300,000 deaths across more than 180 countries **[1]**. The COVID-19 pandemic is a global crisis and control strategies; including the development of PCR-based diagnostics, serological assays, monoclonal antibodies and vaccines, as well as an increased understanding of transmission dynamics, will be informed by knowledge of SARS-CoV-2 sequence data. The sequences in the receptor-binding domain (RBD) of the SARS-CoV-2 viral spike (S) protein, used to bind to cell surface receptors and gain entry into human host cells, are evolutionary close to and possibly spilled over from bat (RaTG13) -like betacoronaviruses, and appear efficient for binding to human ACE2 receptors **[1,2]**. Structural analysis has revealed that the SARS-CoV-2 RBD contains a core and a receptor-binding motif (RBM) that mediates contact with ACE2 **[3]**. Several residue changes in SARS-CoV-2 RBD stabilize two virus-binding hotspots at the RBD and human ACE2 interface and enhance the virus’s binding affinity **[3]**. Therefore, it has been suggested that neutralising monoclonal antibodies targeting SARS-CoV-2 RBM are promising antiviral therapies, and that the RBD itself can function as a subunit vaccine **[3,4]**.

Whilst COVID-19 has spread from China across the globe, the routes of travel are not always clear. Moreover, with this expansion, the likelihood of mutations and viral changes increases, which may impact the efficacy of diagnostics and potentially that of future therapies and vaccines. A growing number of *SARS-CoV-2* whole genomic sequences are available in the public domain **[5,6]** and their phylogenetic analysis can assist researchers and policymakers in answering questions about the spread and evolution of the virus. In general, the phylogeny can be used to identify the transmission network through finding samples with identical sequences, characterising mutations that barcode clades, and inform on underlying host or pathogen phenotypes through genome-wide association methods. In some pathogens, it is possible to detect selection pressure that causes phenotypic traits, through identifying signatures of convergent evolution, that is, where the same causal mutations occur independently in unrelated branches of the phylogenetic tree **[7]**. An increased understanding of the genomic locations that might be under selective pressure could be of value to epidemiologists, disease modellers and vaccine developers. We used more than 5,300 whole genome SARS-CoV-2 virus sequences (size 29.6kbp) in the public domain, sourced globally **[5,6]** (**Table 1**). We explored the mutation diversity across 62 countries to provide insights into the evolutionary history and report any informative mutations for control tools. To assist the ongoing analysis of SARS-CoV-2 virus sequences and to facilitate the investigation of genetic variation, we provide an online tool – “Covid-Profiler” **[8]**.

**Table 1.** Clade frequency (%) across geographic regions.

| Region | No.<br>countries | C1 | C1.1 | C2 | C2.1* | C2.1.1 | C2.1.2 | Overall |
| --- | --- | --- | --- | --- | --- | --- | --- | --- |
| Asia | 22 | 237 (12.1) | 4 (0.2) | 848 (43.4) | 503 (25.7) | 110 (5.6) | 253 (12.9) | 1187 |
| Europe | 26 | 171 (2.2) | 18 (0.2) | 2097 (27.1) | 2134 (27.6) | 1062 (13.7) | 2253 (29.1) | 3682 |
| N. America | 4 | 192 (4.4) | 1073 (24.8) | 408 (9.4) | 271 (6.3) | 2242 (51.9) | 134 (3.1) | 3254 |
| S. America | 7 | 16 (16.3) | 2 (2) | 7 (7.1) | 16 (16.3) | 10 (10.2) | 47 (48) | 2878 |
| Oceania | 2 | 139 (10.9) | 90 (7) | 317 (24.8) | 273 (21.4) | 282 (22.1) | 176 (13.8) | 3725 |
| Africa | 7 | 6 (5.9) | 0 (0) | 5 (4.9) | 57 (55.9) | 19 (18.6) | 15 (14.7) | 761 |
| Total | 68 | 761 (4.9) | 1187 (7.7) | 3682 (23.8) | 3254 (21) | 3725 (24.1) | 2878 (18.6) | 15487 |

\* North America includes one sample from Central America

## Results

An analysis was performed on 15,487 SAR-CoV-2 sequences, sourced from isolates collected between December 24, 2019 and April 29, 2020. A comparison with the Wuhan-Hu-1 reference (NC_045512) resulted in 6,754 unique SNPs in high quality regions, 6,712 (99.4%) of which in genic regions with 4,322 (64.0%) non-synonymous amino acid changes. The number of SNP differences per isolate was small (median 6; range 1 to 17) (**S1 figure**), and the nucleotide diversity across all isolates was low (π = 4.5×10^−7^). A Bayesian SNP-time dated phylogenetic tree (**S2 figure**) highlights the emergence of the virus in China, and its spread through multiple introductions into Europe, North America, and Oceania. It also reveals the clades of transmission involving (near-)identical sequences, initially both within and between countries, but more recently mostly within countries due to nation lockdowns. By looking at the ancestral splits within the tree, there appear to be two main clusters (denoted C1 and C2), which were further partitioned into six main clades (C1, C1.1, C2, C2.1, C2.1.1, and C2.1.2; see **Figure 1a**). Bayesian phylogenetic analysis revealed an association between clade and geographic location, with C2 being more common in all continents (**Figure 1b, Table 1**). The most ancestral cluster was determined to be C1 through phylogenetic reconstruction using the closest known relative to SARS-CoV-2 (MN996532.1). By using ancestral reconstruction of nucleotide states at the most recent common ancestors, we identified thirteen informative barcoding mutations for each clade (**Table 2**). Nine of the clade-defining SNPs are non-synonymous mutations and include D614S in the spike glycoprotein (S) (C2.1) and a triple mutation affecting two codons (R203K, G204R) in the nucleocapsid phosphoprotein (N) (C2.1.2). Interestingly, the three mutations separating C2.1.2 from other isolates are located in the primer binding site of a qPCR kit used by the Chinese CDC **[9]** (**S3 figure**).

**Table 2.** Barcoding SNPs for the six Clades.

| Clade | Genome<br>Position*** | Ancestral/<br>Alternate<br>allele | Amino<br>Acid<br>change | Type | Gene |
| --- | --- | --- | --- | --- | --- |
| C1** | 28144 | ./C* | - | - | *ORF8* |
| C1** | 8782 | ./T* | - | - | *orf1ab:nsp4* |
| C2 | 28144 | C/T | L84S | missense |  |
| C2 | 8782 | T/C | - | synonymous |  |
| C1.1 | 18060 | C/T | S7F | missense | *orf1ab:nsp14* |
| C1.1 | 17858 | A/G | M541V | missense | *orf1ab:nsp13* |
| C1.1 | 17747 | C/T | - | synonymous | *orf1ab:nsp13* |
| C2.1 | 23403 | A/G | D614G | missense | S |
| C2.1 | 14408 | C/T | - | synonymous | *orf1ab:nsp12* |
| C2.1 | 3037 | C/T | - | synonymous | *orf1ab:nsp3* |
| C2.1.1 | 25563 | G/T | Q57H | missense | *ORF3a* |
| C2.1.1 | 1059 | C/T | T85I | missense | *orf1ab:nsp2* |
| C2.1.2 | 28881 | G/A | R203K | missense | N |
| C2.1.2 | 28882 | G/A | R203K | missense | N |
| C2.1.2 | 28883 | G/C | G204R | missense | N |

\* C1 has no ancestral allele as it is the most ancestral type (found using phylogenetic reconstruction using MN996532.1 as an outgroup). S Spike, N nucleocapsid; \*\* match those of L vs. S types in [12]; \*\*\* positions based on Wuhan-Hu-1 reference (NC\_045512)

**Figure 1.**
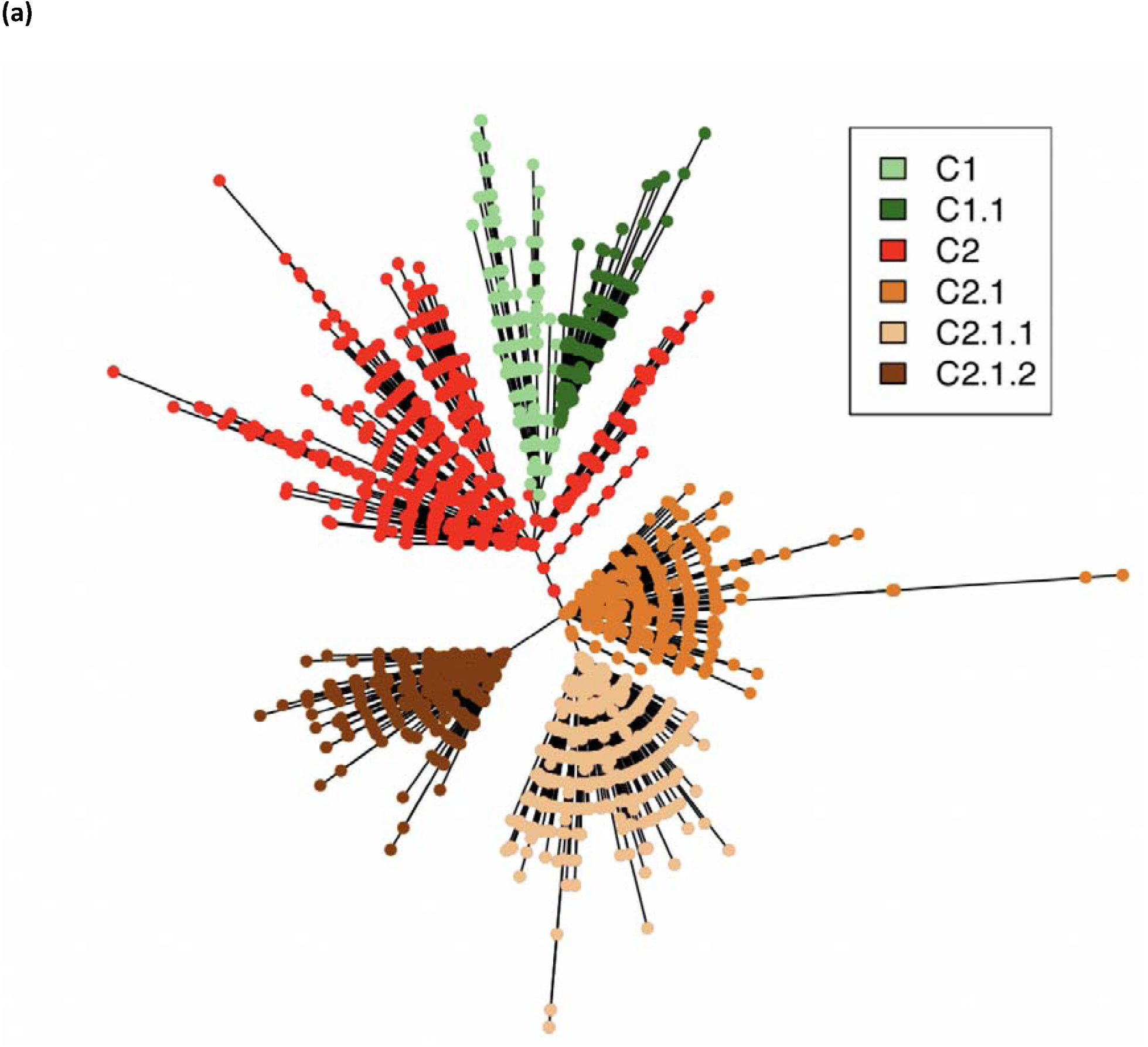
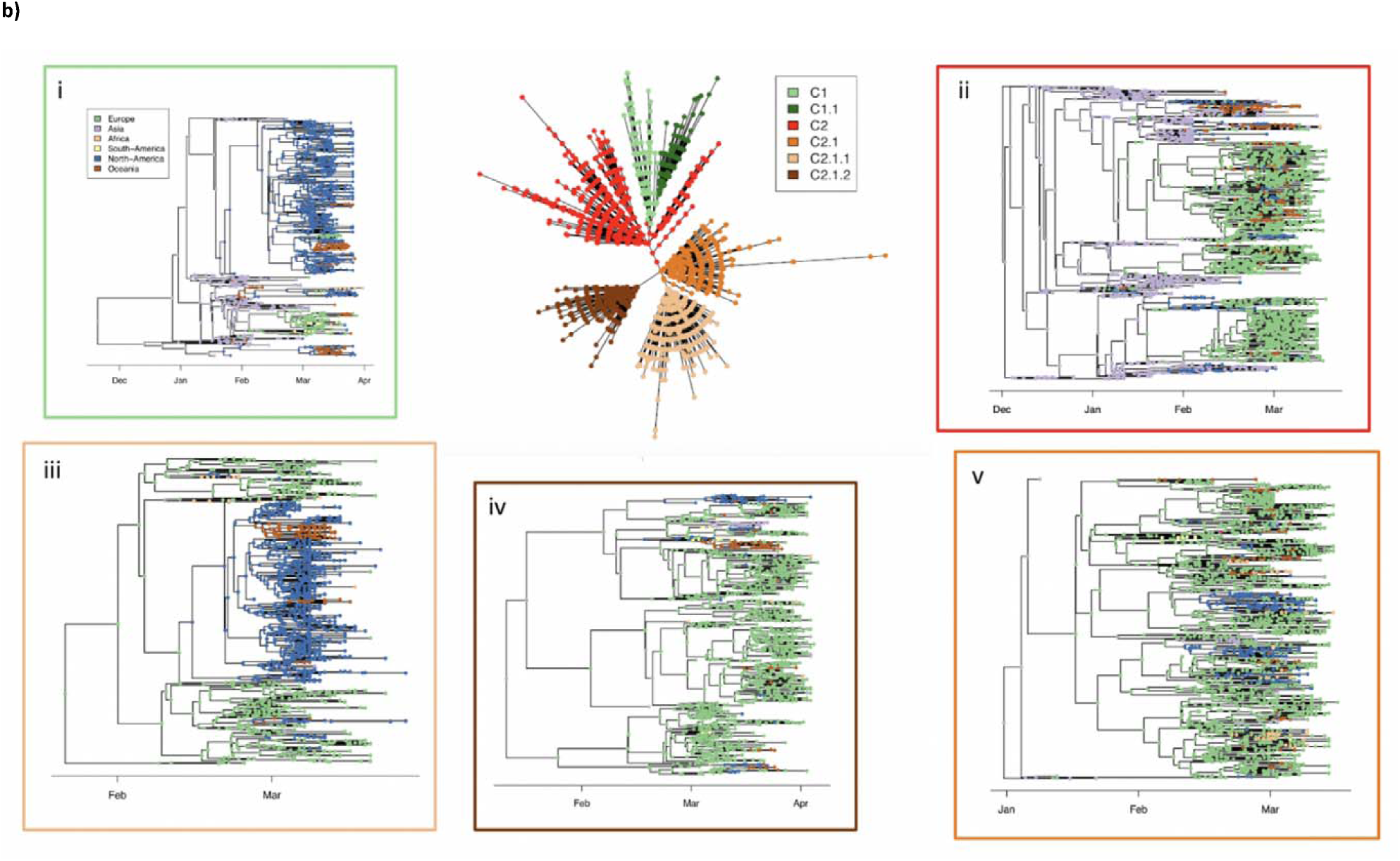
Phylogenetic and clade analysis of 15,487 whole genome SARS-CoV-2 sequences and a BEAST analysis of the first 5,349.

To establish whether particular mutations were under selection we looked for convergent evolution across the whole genome. From a total of 6,294 variant positions, 2,230 sites may have evolved convergently. However, with the relatively large viral genome size and number of isolates, it is expected that random genetic drift will lead to some sites evolving in a convergent manner. An empirical approach was taken to find possible candidates that might be under selective pressure, rather than genetic drift, by looking for outliers from the expected distribution. In particular, using the 99.5 percentile of the distribution of the number of origins, led to a cut-off of 11 independent acquisitions, above which mutations may represent candidates for positive selection. Using this cut-off, and filtering out hits which did not appear in at least 3 countries, we found 14 sites with missense mutations (**Table 3**). The strongest signal was for mutation L37F in *nsp6*, which had 78 independent mutation events. Nsp6 is involved in the formation of double-membrane vesicles (together with *nsp3* and *nsp4*) and is thought to play a role in restricting autophagosome expansion **[10]**. The second strongest hit was the L5F mutation in the spike glycoprotein **(S4 figure)**. These mutations are putatively the result of selection pressure and should be characterized further to understand how the virus is adapting to human transmission.

**Table 3.** Genetic variants with evidence of positive selection by homoplasy.

| Position | No.<br>mutation<br>events* | Ancestral /<br>Alternate<br>allele | Amino<br>acid<br>change | Gene | Number<br>samples<br>** | Countries<br>(Regions<br>***) |
| --- | --- | --- | --- | --- | --- | --- |
| 11083 | 78 | G/T | L37F | *orf1ab:nsp6* | 2049 | 51 (6) |
| 21575 | 41 | C/T | L5F | S | 94 | 18 (4) |
| 3037 | 24 | C/T,A | F106L | *orf1ab:nsp3* | 5604 | 62 (6) |
| 21137 | 17 | A/G | K160R | *orf1ab:nsp16* | 39 | 13 (4) |
| 10323 | 16 | A/G | K90R | *orf1ab:nsp5* | 120 | 9 (4) |
| 19484 | 16 | C/T | A482V | *orf1ab:nsp14* | 23 | 4 (3) |
| 28887 | 16 | C/T | T205I | N | 47 | 12 (4) |
| 1820 | 15 | G/A | G339S | *orf1ab:nsp2* | 28 | 12 (4) |
| 25563 | 15 | G/C,T | Q57H | ORF3a | 3714 | 44 (6) |
| 9474 | 14 | C/T | A307V | *orf1ab:nsp4* | 15 | 6 (3) |
| 14786 | 14 | C/T | A449V | *orf1ab:nsp12* | 135 | 12 (4) |
| 1059 | 13 | C/T | T85I | *orf1ab:nsp2* | 3097 | 36 (6) |
| 9438 | 13 | C/T | T295I | *orf1ab:nsp4* | 21 | 8 (4) |
| 28253 | 13 | C/G,T,A | F120L | ORF8 | 16 | 11 (4) |

S Spike, M Membrane; \* the 99.5 percentile cut-off was 11; \*\* with any alternative allele; \*\*\* from Table 1;

A number of diagnostic primer sets have been defined by the WHO **[9]**. Ideally, primers should target a genomic region which is well conserved among global isolates, or at least so for the geographic catchment area of a diagnostic laboratory. To understand potential differences between primer sets, an analysis of the diversity in their binding sites was performed. For each diagnostic, the number of mismatches in their forward, reverse and probe primer sets was characterised across the 15,487 sequenced isolates (**Table 4**). This revealed that overall diversity was low in binding sites, however, there were some differences between geographic origins. The highest proportion of samples with >=1 mismatch in a primer was the China-N set for which 18.9% of global isolates had a mutation. When analysed on the country level there were marked differences with some countries in Latin America and Africa showing >70% of samples carrying a mutation in the primer binding site (**S3 figure**).

**Table 4.**
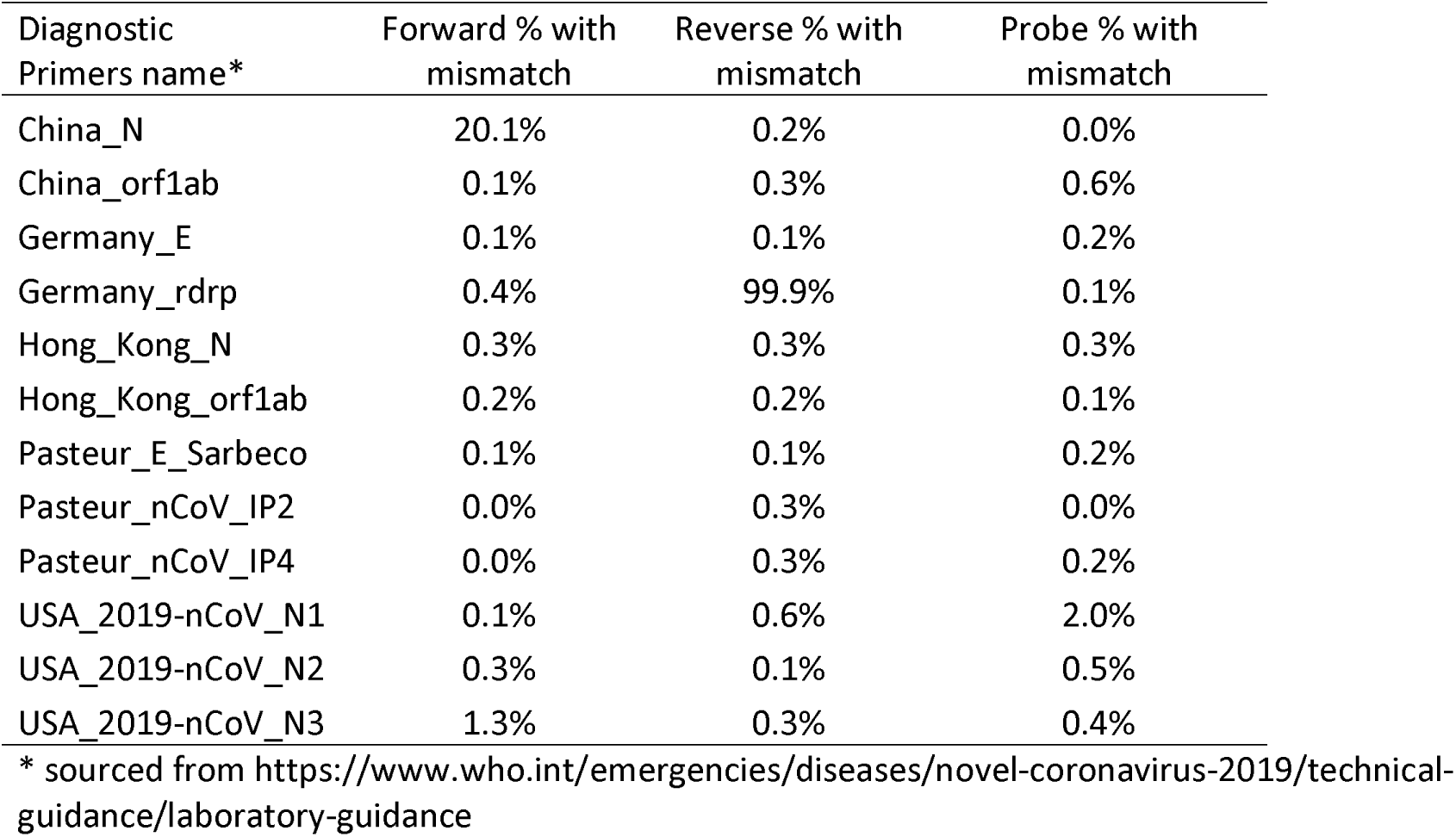
Reported COVID-19 PCR diagnostic assays, with their associated frequency of mismatches identified across the 5,349 SARS-CoV-2 sequences

| Diagnostic Primers name* | Forward % with mismatch | Reverse % with mismatch | Probe % with mismatch |
| --- | --- | --- | --- |
| China_N | 20.1% | 0.2% | 0.0% |
| China_orf1ab | 0.1% | 0.3% | 0.6% |
| Germany_E | 0.1% | 0.1% | 0.2% |
| Germany_rdrp | 0.4% | 99.9% | 0.1% |
| Hong_Kong_N | 0.3% | 0.3% | 0.3% |
| Hong_Kong_orf1ab | 0.2% | 0.2% | 0.1% |
| Pasteur_E_Sarbeco | 0.1% | 0.1% | 0.2% |
| Pasteur_nCoV_IP2 | 0.0% | 0.3% | 0.0% |
| Pasteur_nCoV_IP4 | 0.0% | 0.3% | 0.2% |
| USA_2019-nCoV_N1 | 0.1% | 0.6% | 2.0% |
| USA_2019-nCoV_N2 | 0.3% | 0.1% | 0.5% |
| USA_2019-nCoV_N3 | 1.3% | 0.3% | 0.4% |

\* sourced from https://www.who.int/emergencies/diseases/novel-coronavirus-2019/technical-guidance/laboratory-guidance

The spike glycoprotein mediates cell entry by binding to host ACE2 and has been proposed as a target for antibody-mediated neutralisation **[11]**. It is important to take into account the variation in the amino acid sequence as this could influence the efficacy of the vaccine. Across the 15,487 SAR-CoV-2, we found 841 high quality mutations in the spike gene, with 543 missense and 298 synonymous mutations. The mutations were generally rare, with 54.6% occurring in only single isolates. One high-frequency spike mutation, D614G, was acquired on the branch defining C2.1, leading to just over 63.8% of all isolates containing this mutation (**Figure 3A**). The SNPs were spread throughout the gene, with a sliding window analysis of the SNP density (using windows of 50nt) reporting a generally even distribution (range: 1 to 22 SNP positions, **Figure 3A**). Analysis of the protein structural model indicated some of these mutations present on the interface between spike and ACE2 (**Figure 3B, S4 Figure**) and could potentially affect vaccine performance if selection pressure (such as vaccine escape) drove them to higher frequencies. Although these mutations are currently rare, they need to be evaluated in case they affect vaccine performance.

**Figure 2.**
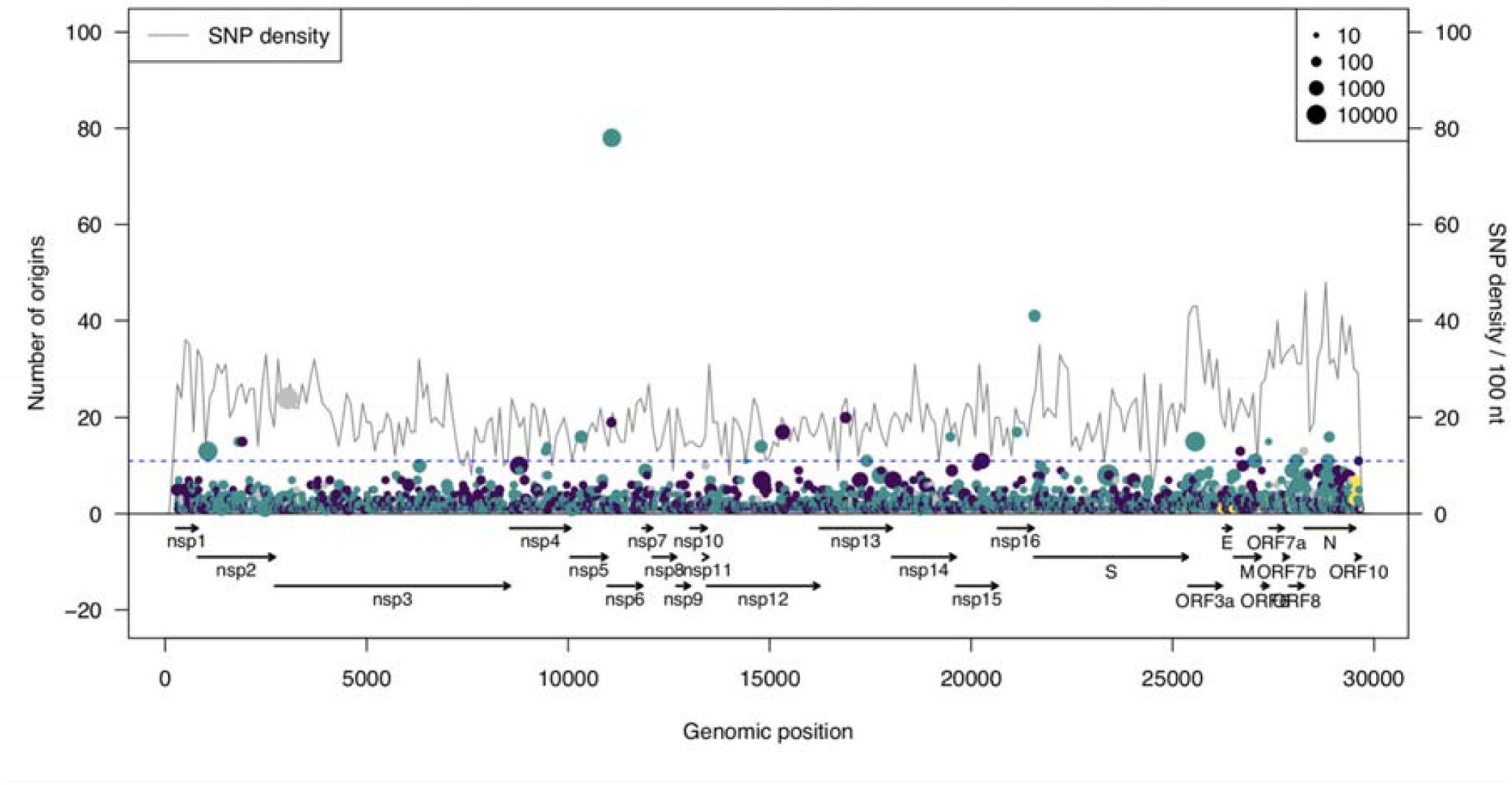
The genetic diversity across the *SARS-CoV-2* genome. Each SNP is represented by a circle. The number of times the SNP has been independently acquired is indicated on the Y-axis. The size of the circle represents the derived allele frequency. The circles are coloured to represent intergenic (yellow), synonymous (purple), missense (green) or a mix of the aforementioned (grey). The line plot represents the SNP density across the genome in windows of 100 nucleotides.

**Figure 3.**
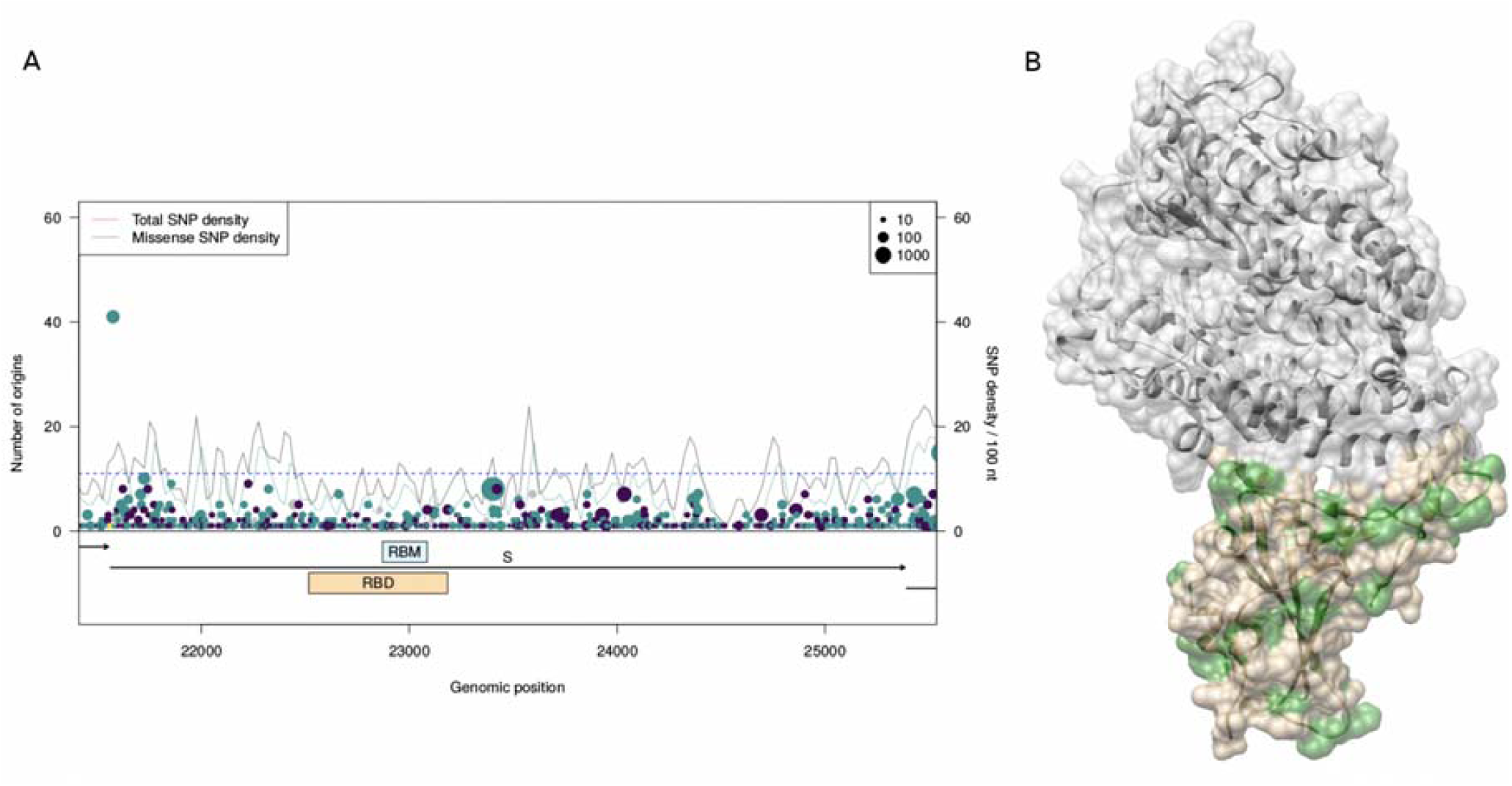
The SNPs in the spike glycoprotein (S) A) A gene level view of genetic variation in S is shown, with each SNP represented by a circle. The number of times the SNP has been independently acquired is indicated on the left Y-axis. The size of the circle represents the derived allele frequency. The circles are coloured to represent synonymous (purple), missense (green) or a mix of the aforementioned (grey). The line plots represents the SNP density, total (grey) and non-synonymous (green), across the gene in windows of 100 nucleotides (Y-axis right hand scale). The location of the receptor binding domain (RBD, in brown) and the receptor binding motif (RBM, in blue) are highlighted on the gene track. B) The structure of the receptor binding domain of S (brown) bound to ACE2 (grey). Mutation positions in S are highlighted in green.

## Discussion

Our analysis, based on 15,487 SAR-CoV-2 genomes, shows a clear segregation into clades of transmission within and between countries. We report two main clades which are further clustered into six sub-clades, defined by thirteen informative barcoding mutations. There is overlap between our two main clades (C1 vs. non-C1) and the L and S types defined using only 103 isolates [**12**], where the two separating or barcoding mutations (in *orf1ab* and *ORF8)* are identical. Our phylogenetic analysis provides much finer-scale resolution on the phylogeny of C2 into sub-clades, especially in the context that SARS-CoV-2 has been accumulating an average of one to two mutations per month. Having arisen recently, the level of diversity observed are much lower than for other viruses (e.g. dengue) where subtypes or lineages exist with many more SNP differences and associated functional differences [**13**]. Whilst functional follow-up is required for the barcoding mutations, our SNPs (or updated versions) are likely to be useful once countries have attained greater control over domestic transmission chains, where the ability to understand and map the geographic sources of viral importation might become crucial for effective disease control.

Our analysis furthermore reveals fourteen non-synonymous sites that putatively might be subject to convergent evolution. Of these, the two strongest SNPs are located in *nsp6 and* the spike glycoprotein. Further research might be warranted to elucidate the role of these mutations and to further characterize the evolutionary benefits encoded by these mutations. In the future, when effective antivirals may have been developed and deployed, the tracking of the occurrence of convergent SNPs might further gain in importance and utility in monitoring the onset of antiviral resistance.

The limitations of our study are the relatively small proportion of sampled sequences (>15,000) compared to infections (>4,000,000), an over-reliance on sequences from relatively severe cases, and a potential bias towards collecting more isolates early in an outbreak when most genomes are still very similar. Similarly, much of the sequencing to date has been performed in the USA, UK and Australia. If not accounted for, this bias can lead to false inferences on the impact of particular mutations on transmission. One such example is the D614G mutation [14] which was proposed as leading to strains becoming more transmissible, but further work has identified as a homoplasy site not associated with transmission [15]. Our work has shown this site to be introduced in the C2.1 clade and all subclades, and although it does occur in other clades due to homoplasy, it fails to reach our significance cutoff, with the bulk of the isolates harbouring this mutation arising from a single mutation event in a founder strain. The resulting bias in cluster size means we cannot infer that larger clades are more virulent or more transmissible. However, we could use the transmissibility phenotype, or others involving laboratory-based virulence or patient disease severity, to identify related causal mutations using GWAS or convergent evolution techniques [16]. One roadblock is the small number of polymorphisms observed in SARS-CoV-2, and the unknown contributing role of host genetics and immune system. Understanding the role of host genetics in host-virus interactions with regards to pathogenesis and disease progression is also important and await studies of sufficient size to be conclusive.

This study has also highlighted a number of SNPs that might cause misalignment with commonly used diagnostic PCR primer sets and might thus hamper screening programs and patient diagnosis through providing false negative test results. These mismatches were especially common in Latin America. Our “Covid-profiler” tool allows for the upload of SARS-CoV-2 sequence data to infer informative mutations, update phylogenies, and assess the robustness of diagnostics, as well as other control tools like vaccines. As SARS-CoV-2 virus is likely to become the fifth endemic coronavirus in the human population, and probably will not be the last coronavirus to jump the animal-to-human species boundary, our “Covid-profiler” is likely to have utility beyond the current outbreak.

## Conclusions

Analysis of global SARS-CoV-2 viral sequences has led to insights into the robustness of existing, and development of new, tools for infection control. We have developed an online tool (“Covid-profiler”) to facilitate this type of analysis and profile SARS-CoV-2 viral sequences.

## Methods

Full SARS-CoV-2 genome sequences were downloaded from the GISAID [6] and NCBI [5], covering isolates collected between December 24, 2019 and April 6, 2020. Sequences were aligned to the reference genome (NC_045512.2) using mafft software [**17**]. IQ-TREE (v1.6.12) [**18**] and BEAST (v1.10.4) [**19**] software were used to reconstruct the phylogeny tree. Time dated trees were generated by BEAST software using an HKY site model with 4 gamma categories, the coalescent exponential model and an uncorrelated relaxed clock rate. The number of independent acquisitions of the mutation was counted using empirical Bayesian ancestral reconstruction methods and custom scripts utilising the ete3 python library **[20]**. A high number of acquisitions can be indicative of genomic regions under positive selection pressure. The SARS-CoV-2 spike glycoprotein structures was downloaded from the PDB (6m0J and 6vxx) **[21, 22]** and visualised using chimera software **[23]**.

## DECLARATIONS

### Ethics approval and consent to participate

Not applicable

### Consent for publication

Not applicable

### Availability of data and materials

The datasets analysed during the current study are available from GISAID (https://www.gisaid.org) and NCBI (https://www.ncbi.nlm.nih.gov)

### Authors’ contributions

JP, SC, MLH, and TGC conceived and directed the project. JP, WD and DW coordinated data processing. JP and WD performed bioinformatic and statistical analyses under the supervision of MLH and TGC. All authors interpreted results. JP, WD, MLH and TGC wrote the first draft of the manuscript. All authors commented and edited on various versions of the draft manuscript and approved the final manuscript. JP compiled the final manuscript.

## Acknowledgements

We thank Aleksei Ponomarev for providing coding support on the data processing. We gratefully acknowledge the laboratories who submitted the sequences to the NCBI and GISAID public databases on which this research is based. We also thank NCBI and GISAID for developing and curating their databases.

---

**S1 figure.**
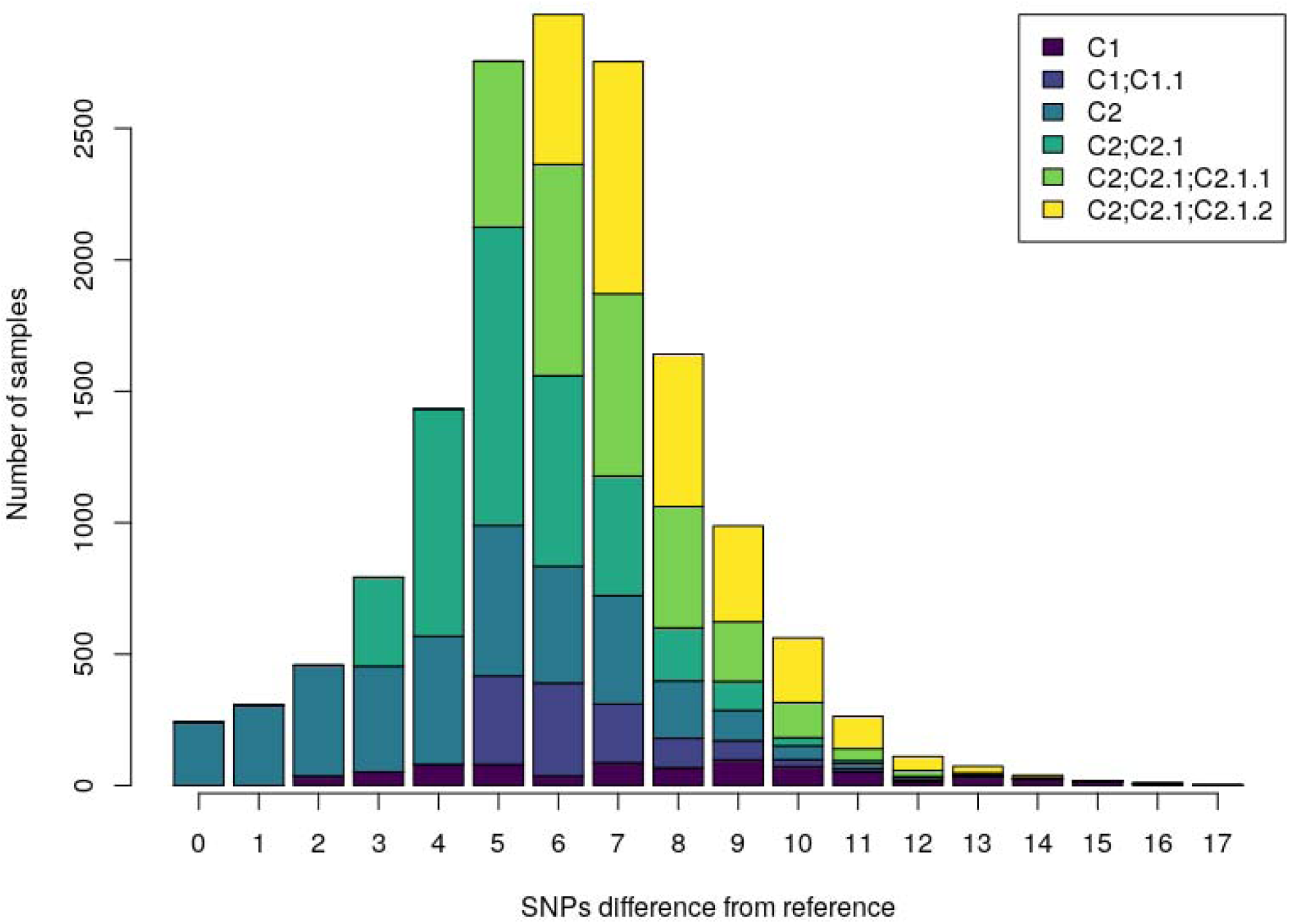
Distribution of SNP differences from the Wuhan-Hu-1 reference (NC_045512) by clade

**S2 figure.**
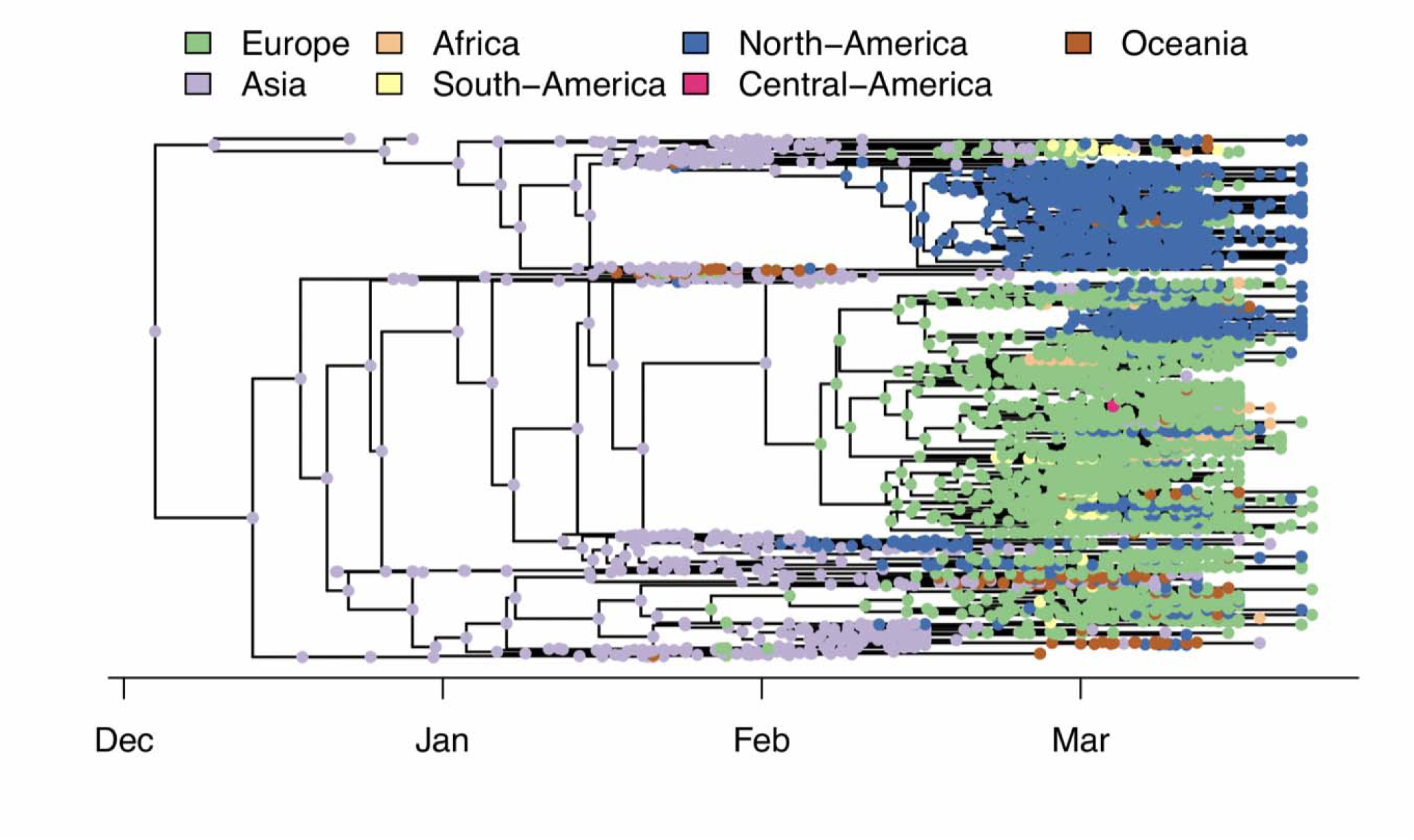
Bayesian time-dated tree of the first 1,200 SARS-CoV-2 genomes, demonstrating origin of the clades. Ancestral reconstruction of the continent of isolation confirms known patterns of spread throughout the globe

**S3 figure.**
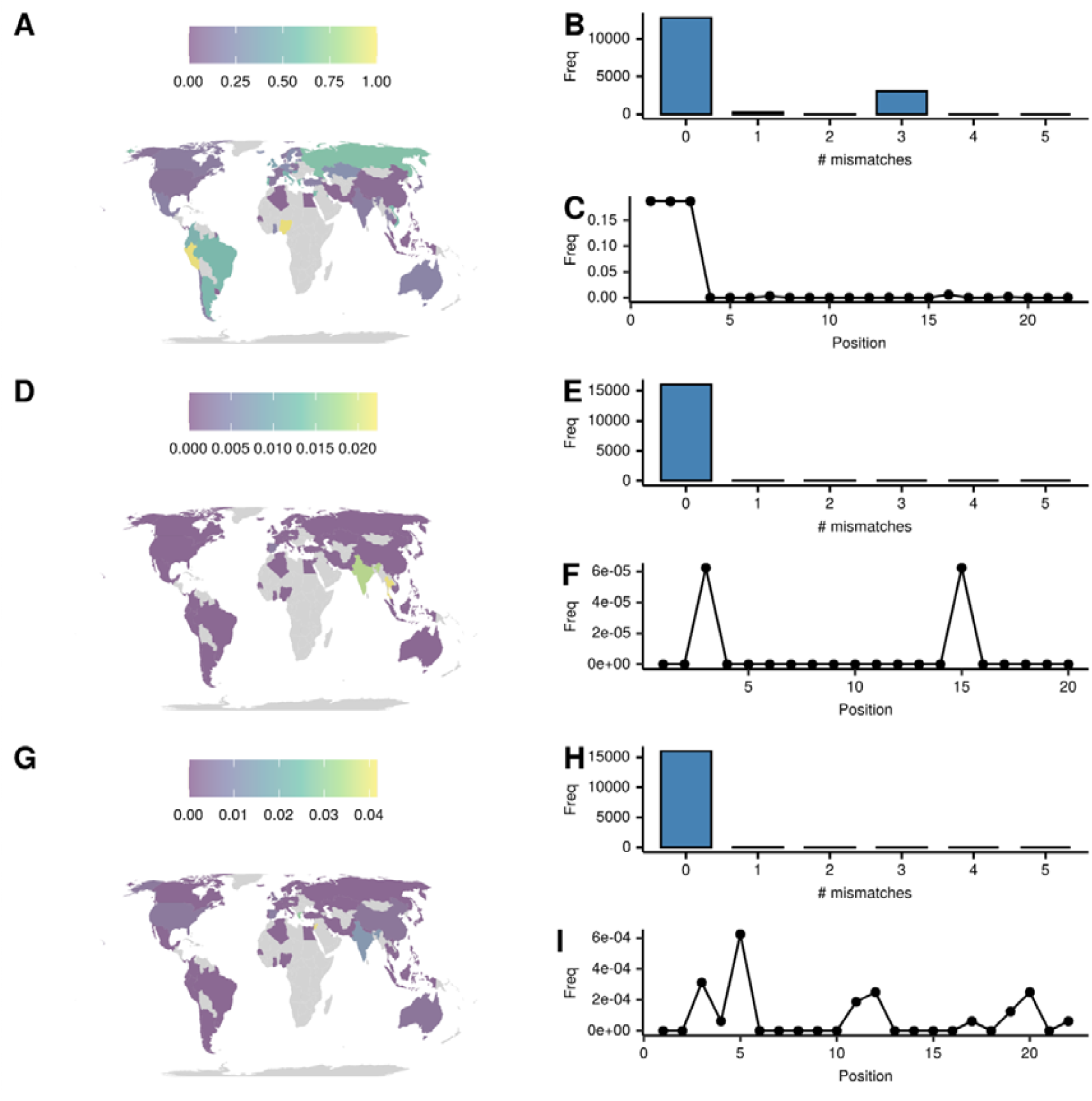
The global distribution of the mutation pattern for the PCR diagnostic primers and prob of the China-N assay. Panel A shows the fraction of isolates with at least one mutation in the forward primer in different countries. Panel B shows the number of samples with different numbers of mismatches. Panel C shows the frequency and location of mismatches across the primer binding site. Panels D, E and F show the same metrics for the probe and panels G, H and I for the reverse primer.

**S4 figure.**
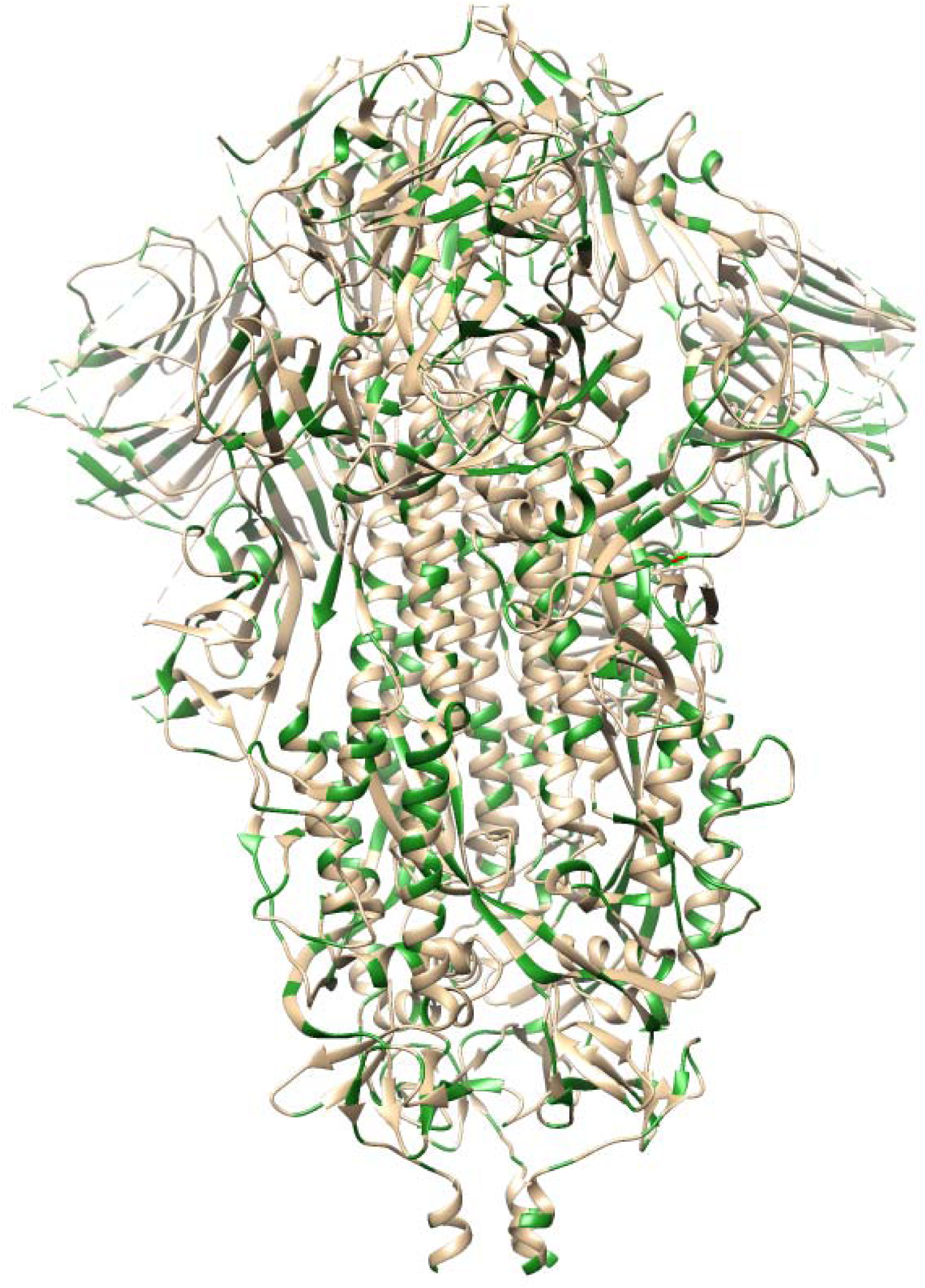
Mutations visualised on the spike glycoprotein. SNP mutations are presented in green. The C2.1 defining SNP D614G is highlighted in red

